# Isoprenoid Binding and Substrate Channeling in Drimenol Synthase, a Bifunctional Class II Terpene Cyclase-Phosphatase

**DOI:** 10.64898/2026.08.29.748031

**Authors:** Kristin R. Osika, Mason E. Leffler, Briana Abigail R. Czarnecki, David W. Christianson

## Abstract

More than one thousand bifunctional terpene synthases combining prenyltransferase and terpene cyclase activities have been identified to date, but only a handful of enzymes have been identified that combine terpene cyclase activity with a downstream processing activity. Drimenol synthase from the marine bacterium *Aquimarina spongiae* (AsDMS) consists of a class II terpene cyclase that converts farnesyl diphosphate into drimenyl diphosphate, and a haloacid dehalogenase-like phosphatase that hydrolyzes drimenyl diphosphate to generate the sesquiterpene alcohol drimenol. The first crystal structure of AsDMS to be reported revealed the architecture of domain assembly as well as dimeric quaternary structure, establishing a structural chemical foundation for cyclization and hydrolysis mechanisms [K. R. Osika, M. N. Gaynes, D. W. Christianson (2025) *Proc. Natl. Acad. Sci. U.S.A. 122*, e2506584122]. Here, we report crystal structures of the catalytically-inactive double mutant, D33A-D323A AsDMS, complexed with farnesyl diphosphate, geranyl diphosphate, and dimethylallyl diphosphate. The diphosphate group of each isoprenoid dominates binding interactions in the active sites of both the cyclase and phosphatase domains. In the cyclase active site, only farnesyl diphosphate is sufficiently long for its terminal isoprenoid C=C bond to bind adjacent to the catalytic general acid that would initiate the cyclization cascade in the wild-type enzyme. In the phosphatase active site, all isoprenoid diphosphate groups bind similarly, but isoprenoid chain conformations vary. These structures provide a foundation for understanding substrate recognition and catalysis in both active sites. Finally, we present kinetic evidence suggesting that substrate channeling may be operative in wild-type AsDMS.

## Introduction

Terpenoids represent the largest and most structurally diverse family of natural products, encompassing more than 100,000 known compounds distributed across all domains of life.^1,2^ These molecules fulfill important ecological functions in plants, fungi, bacteria, and animals, and many are commercially valuable due to a wide variety of applications in medicine, agriculture, and industry.^3–7^ All terpenoids derive from two universal C_5_ precursors, dimethylallyl diphosphate (DMAPP) and isopentenyl diphosphate (IPP), which are linked together by a prenyltransferase to generate longer isoprenoids such as C_10_ geranyl diphosphate (GPP), C_15_ farnesyl diphosphate (FPP), or C_20_ geranylgeranyl diphosphate (GGPP).^8–10^ In turn, a linear isoprenoid such as FPP is converted into a complex, multi-ringed product through a carbocation-mediated cyclization cascade catalyzed by a terpene cyclase.^11–14^

Terpene cyclases are mainly grouped into two classes based on their domain architecture and first mechanistic step^13,15^ (some noncanonical cyclases have also been identified^16^). Class I cyclases exhibit α, αβ, or αβγ domain architecture and initiate cyclization reactions through the metal-dependent ionization of the substrate diphosphate group in the α domain. Class II cyclases exhibit β, βγ, or αβγ domain architecture and initiate cyclization reactions in the β domain through protonation of the terminal isoprenoid C=C bond by an aspartic acid general acid.^13,17,18^

Some terpene synthases are bifunctional^19,20^ – most common are enzymes that combine prenyltransferase and cyclase activities, such as fusicoccadiene synthase.^21–24^ Others catalyze tandem cyclization reactions, such as geosmin synthase^25,26^ or abietadiene synthase.^27–29^ Less common are bifunctional enzymes that combine a terpene cyclase with a downstream tailoring activity. For example, drimenol synthase from the marine bacterium *Aquimarina spongiae* (AsDMS, UniProt A0A1M6CXF0) consists of a class II cyclase in a single β domain that converts FPP into drimenyl diphosphate (DPP), and a haloacid dehalogenase (HAD)-like phosphatase that hydrolyzes DPP to form drimenol (Figure 1).^30^ We recently reported the first crystal structure of AsDMS, revealing that the catalytic domains of the enzyme assemble such that two distinct active sites are positioned on opposite sides of the protein.^31^ Our accompanying mechanistic studies further established that the hydroxyl oxygen of drimenol derives from the prenyl oxygen of FPP and not from bulk water; we also established that the hydrolysis of DPP occurs in stepwise fashion to yield drimenol plus two equivalents of inorganic phosphate (P_i_). These structural and enzymological studies pointed to a unified chemical mechanism for tandem class II cyclization-dephosphorylation reactions.^31^

**Figure 1.**
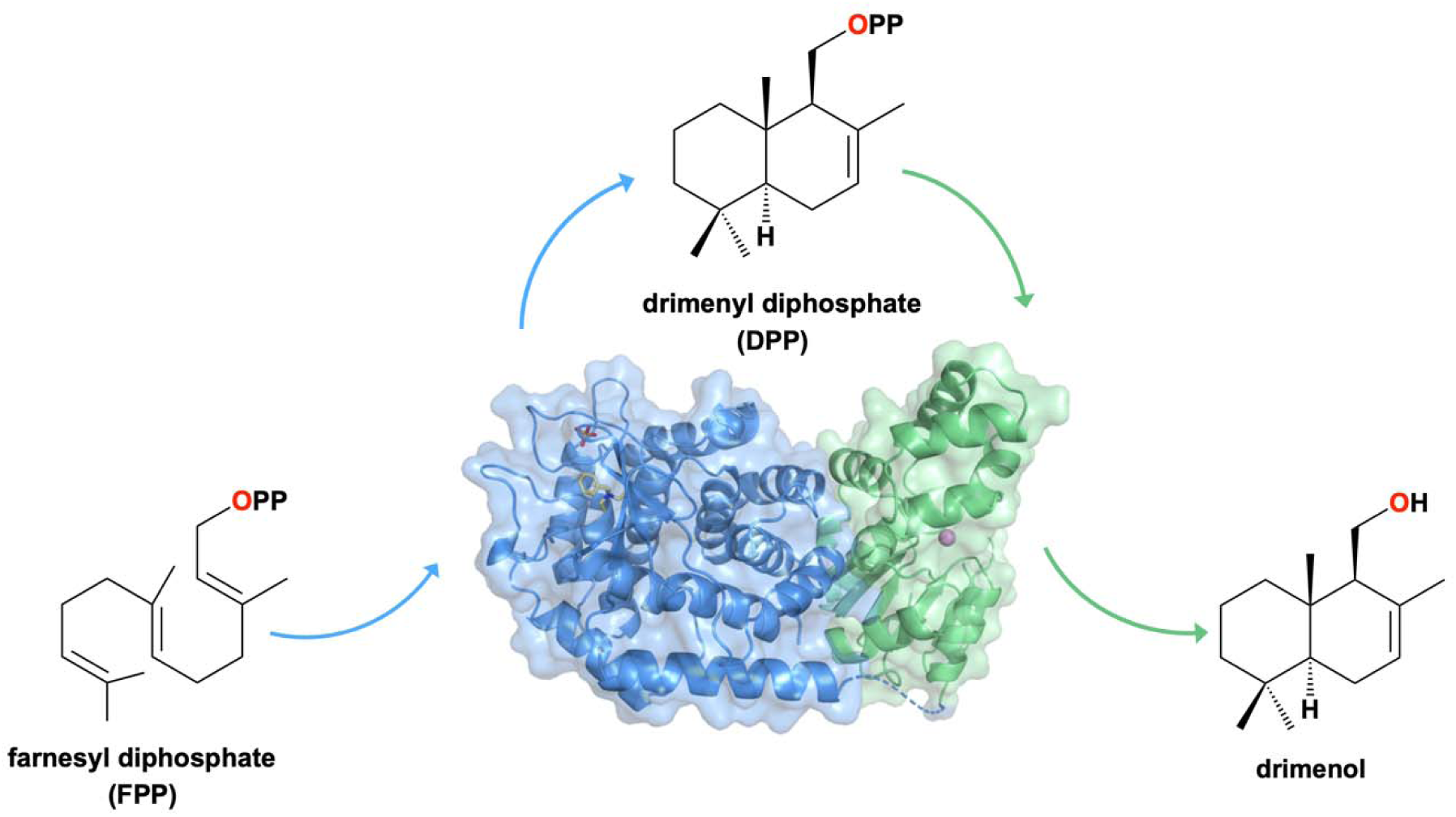
AsDMS structure (PDB 9MHS) and reaction sequence. Class II terpene cyclase β domain (blue) and HAD-like phosphatase domain (green) active sites are located on opposite sides of the protein structure. Intermediate DPP transits from the cyclase to the phosphatase for stepwise hydrolysis of the diphosphate group to yield drimenol. The prenyl oxygen of FPP is retained as the hydroxyl oxygen of drimenol.

Takahashi and colleagues subsequently and independently reported the AsDMS structure, confirming our previously reported structure and unified chemical mechanism.^32^ These investigators also reported the 2.3 Å resolution crystal structure of D333N AsDMS complexed with the intact substrate FPP, which binds in the precise conformation that would be required to generate the *trans*-decalin skeleton of drimenol. Substitution of the catalytically obligatory general acid in the β domain abolished catalytic activity and enabled stabilization and crystallization of the precatalytic enzyme-substrate complex.

Here, we further explore isoprenoid binding in both catalytic domains of AsDMS through the study of a double mutant lacking the general acid in the class II cyclase β domain as well as the catalytic nucleophile/Mg^2+^ ligand in the phosphatase domain, D33A-D323A AsDMS (due to different numbering conventions, D323 in our construct is equivalent to D333 in the Takahashi construct^32^). Cocrystallization of D33A-D323A AsDMS with FPP, GPP, and DMAPP yields structures showing intact isoprenoid diphosphates bound in both active sites. Additionally, the reaction kinetics of wild-type AsDMS compared with the reaction kinetics of an equimolar mixture of single-point mutants D33A AsDMS and D323A AsDMS reveal a catalytic advantage for catalysis by the wild-type enzyme, suggesting that some of the DPP intermediate can transit from the class II cyclase β domain to the phosphatase domain without release to bulk solvent. In other words, intramolecular substrate channeling may be operative in wild-type AsDMS.

## Materials and Methods

### Expression and purification

The previously reported^31^ pET-His₆-MBP-TEV-AsDMS plasmid lacking the first 10 residues at the N-terminus was used as a starting point for designing the D33A-D323A double mutant; additionally, the maltose binding protein (MBP) solubility tag was replaced by green fluorescent protein (GFP). The resulting pET-His₆-GFP-TEV-AsDMS(D33A-D323A) plasmid was codon-optimized for expression in *Escherichia coli* and obtained from GenScript (Piscataway, NJ). Expression and purification of this construct were carried out as described previously.^31^ The pET-His₆-GFP-TEV-AsDMS plasmids of D33A (phosphatase inactive) and D323A (cyclase inactive) AsDMS similarly lacking the first 10 residues at the N-terminus were transformed into DE3 BL21 competent *E. coli* cells (New England Biolabs) and grown at 37° C overnight on Luria-Bertani (LB) agar plates supplemented with 50 µg/mL kanamycin. Colonies were inoculated into 300 mL LB media with 50 µg/mL kanamycin. After overnight incubation and orbital shaking at 37° C and 230 rpm, 30 mL of each starter culture was added to 6×2-L baffled flasks containing 1 L of LB broth and 50 µg/mL kanamycin. These 1-L cultures underwent orbital shaking at 37° C and 230 rpm until reaching an optical density (OD_600_) of 0.6-0.8. Expression of the protein was induced by adding isopropyl-β-D-thiogalactopyranoside (IPTG) to a final concentration of 1 mM in each 1-L culture and incubated overnight in the orbital shaker at 18° C and 180 rpm. Cells were pelleted via centrifugation (20 min, 6,000 rpm, 4° C) and the pellet was collected and resuspended in 50 mM NaH_2_PO_4_•H_2_O (pH 7.3), 500 mM NaCl, 20% glycerol (lysis buffer).

Each suspension was treated with 100 mg lysozyme (GoldBio), 10 mg DNase I (Roche), and one EDTA-free cOmplete Mini Protease Inhibitor Tablet (Roche), left to stir at room temperature for 1 h. After suspension, the cells underwent lysis using a Q700 sonicator (QSonica) at 30% amplitude for 10 min (1 sec on, 2 sec off). The lysate was clarified by centrifugation at 18,000 rpm for 40 min at 4° C. The supernatant was loaded onto a 5-mL HisTrap column preequilibrated with lysis buffer (GE Healthcare) at 3 mL/min and eluted with a 50 mL gradient to 100% 50 mM NaH_2_PO_4_•H_2_O (pH 7.3), 500 mM NaCl, 20% glycerol, 200 mM imidazole (elution buffer). Fractions were analyzed using sodium dodecyl sulfate-polyacrylamide gel electrophoresis (SDS-PAGE), and fractions containing protein of the appropriate molecular weight were collected and treated with 6 mg TEV protease while undergoing dialysis in lysis buffer overnight at 4° C. Tag cleavage was determined to be complete and comparable across all preparations by SDS-PAGE. This sample was then loaded onto a 5-mL HisTrap column, and the flowthrough was analyzed by SDS-PAGE. Fractions containing the protein of interest were pooled and concentrated to approximately 15-21 mg/mL.

### Enzyme product assays

To detect and quantify any possible alcohol products generated by wild-type AsDMS and the D33A, D323A, and D33A-D323A mutants, gas chromatography–mass spectrometry (GC–MS) was employed using an Agilent 8890 GC system coupled to a 5597C MSD and equipped with a J&W HP-5 ms Ultra Inert capillary column (30 m × 0.25 mm × 0.25 μm). Assays were performed in triplicate with a 200-μL total volume, with 5 μM enzyme in reaction buffer [20 mM Tris-HCl (pH 7.3), 150 mM NaCl, 2 mM MgCl₂·6H₂O, 1 mM TCEP, and 10% glycerol]. Stock solutions of 10 mM DMAPP, GPP, FPP, and GGPP (Isoprenoids.com) were made by resuspension in 7:3 methanol/10 mM NH_4_HCO_3_. A 10 mM stock of farnesol (Fisher) was made by resuspension in ethanol. Reactions were initiated upon the addition of 10 μL of 10 mM substrate stock solution, followed immediately by an overlay of 200 μL hexanes containing a 750 μM hexadecane internal standard. Each reaction was incubated at room temperature overnight (15 h), then quenched by vortexing (10 s) and centrifugation (15 s at 12,000 rpm). A 100 μL aliquot of the organic phase was collected for GC-MS analysis. For all samples, the temperature program was initiated with an oven temperature of 60 °C sustained for 2 min, followed by a ramp of 20 °C/min to reach 240 °C. MS data were acquired in positive electron ionization (EI) mode after a 3-min solvent delay. Products were characterized through comparison of mass spectra and chromatograms to data in the National Institute of Standards and Technology database.

### Mass photometry

Experiments were performed at the Johnson Foundation Structural Biology and Biophysics Core at the Perelman School of Medicine, University of Pennsylvania (Philadelphia, PA). Briefly, mass photometry measurements utilized a Refeyn 2MP mass photometer, MassGlass UC cover slips, and silicone gaskets. Droplet dilution was performed by adding 15 μL buffer [150 mM NaCl, 20 mM Tris (pH 7.3), 1 mM TCEP] to the sample well, followed by focusing. Enzyme (5 μL) was then added to reach target final concentrations. Optical contrast was converted into molecular weight using a bovine serum albumin and bovine thyroglobulin calibration standard diluted into the same buffer conditions. A 60-s movie was recorded, and 3250 landing events were counted with AcquireMP software. Analysis and histogram generation were performed with DiscoverMP software.

### Substrate Channeling

To test for substrate channeling, the EnzChek Phosphate Assay (Fisher) was used to quantify the amount of inorganic phosphate released during the reaction. Reactions totaling 200 µL containing equimolar wild-type AsDMS or D33A AsDMS + D323A AsDMS at various concentrations in reaction buffer [20 mM Tris-HCl (pH 7.3), 150 mM NaCl, 1 mM TCEP, 2 mM MgCl_2_, 0.25 mM 2-amino-6-mercapto-7-methylpurine riboside (MESG), 1x Enzchek Reaction Buffer, and 0.2 U purine nucleotide phosphorylase (PNP)] were pre-incubated for 10 min at room temperature to quench any pre-existing phosphate. Varying concentrations (0–250 µM) of FPP were added to initiate reactions, which were monitored in experimental triplicate at 360 nm in a kinetic loop using a Spark Microplate Reader (Tecan) for 45 min at room temperature. PNP phosphorylates the ribose ring of MESG in the presence of inorganic phosphate, affording 2-amino-6-mercapto-7-methylpurine, which has a strong absorbance at 360 nm. The change in A_360_ thus indicates the production of inorganic phosphate. After subtraction of baseline absorbance (0 µM FPP), phosphate concentrations and product concentrations were derived by comparison to a standard curve.

Linear regression was performed using the product concentrations as a function of time. The initial velocity (v_0_) corresponded to the slope of the fitted line, and the uncertainty in v_0_ was the standard deviation calculated for the three experimental replicates.

### Crystal Structure Determinations

The sitting-drop vapor diffusion method was used to crystallize D33A-D323A AsDMS complexes with FPP, GPP, and DMAPP using a Mosquito crystallization robot (SPT Labtech). For structures in complex with FPP or GPP, a 100-nL drop of protein solution [12 mg/mL D33A-D323A AsDMS, 20 mM Tris-HCl (pH 7.3), 150 mM NaCl, 1 mM TCEP, 10% glycerol, 2 mM FPP or GPP, 2 mM tungstate, 2 mM MgCl_2_·6H_2_O] was added to a 100-nL drop of precipitant solution [0.1 M sodium citrate tribasic dihydrate (pH 5.0), 30% Jeffamine ED-2001 (pH 7.0), 0.1 M yttrium (III) chloride hexahydrate] and equilibrated against a 50-µL reservoir of precipitant solution. For the structure in complex with DMAPP, a 100-nL drop of protein solution [7 mg/mL D33A-D323A AsDMS, 20 mM Tris-HCl (pH 7.3), 150 mM NaCl, 1 mM TCEP, 10% glycerol, 2 mM DMAPP, 2 mM MgCl_2_·6H_2_O] was added to a 100-nL drop of precipitant solution [0.1 M sodium citrate tribasic dihydrate (pH 5.5), 26% Jeffamine (pH 7.0)] and equilibrated against a 50-µL reservoir of precipitant solution. Crystals generally appeared within 7 days; prior to flash-cooling in liquid nitrogen, crystals were cryoprotected by submersion in mother liquor augmented with 20% (v/v) glycerol.

All X-ray diffraction data were collected at the 17-ID-2 FMX beamline at the National Synchrotron Light Source II (NSLS-II), Brookhaven National Laboratory (Upton, NY). Diffraction data were indexed, integrated, and scaled with XDS^33^ and merged with AIMLESS as part of the autoPROC pipeline.^34^ Molecular replacement with the PHASER module of PHENIX^35,36^ was used to phase each initial electron density map with the atomic coordinates of the AsDMS-BTAC-P_i_-Mg^2+^ complex (PDB 9MHS) used as a search probe for rotation and translation functions. Each initial atomic model underwent a round of manual model building using Coot,^37^ followed by iterative rounds of manual model building and crystallographic refinement using Coot and PHENIX. During model building, side chains exhibiting no or poor electron density in a 2|F|_o_ – |F|_c_ map contoured at 1.5σ were deleted. Additionally, some polypeptide segments were disordered and thus excluded from the final model. In monomer A of the FPP complex, these segments included residues 1–10 of the construct used for crystallization (corresponding to residues 11–20 of the full-length protein) and residues 215–225 (the interdomain linker). In monomer B of the FPP complex, disordered segments included residues 1–11 and 216–223. In monomer A of the GPP complex, these segments included residues 1–10, 215–225, and 298– 302; in monomer B, disordered segments included residues 1–11 and 215–225. In monomer A of the DMAPP complex, disordered segments included residues 1–12, 22–25, and 215–225; in monomer B, these segments included residues 1–11, 217–225, and 298–302. FPP, GPP, and DMAPP were built into their respective electron density maps in the final stages of refinement. In the cyclase domain of monomer A in the DMAPP complex, additional electron density trailing from the vicinal dimethyl group of DMAPP was observed and remained unmodeled, perhaps arising from the binding of a PEG or Jeffamine fragment. DMAPP was not modeled in the cyclase domain of monomer B as the density was weak and discontinuous. MolProbity was used to validate final refined structures.^38^ Data collection and refinement statistics are summarized in Table 1.

**Table 1.** Data collection and refinement statistics for D33A-D323A AsDMS complexes.

| Complex | FPP | GPP | DMAPP |
|---|---|---|---|
| Space group | *P*4<sub>1</sub>2<sub>1</sub>2 | *P*4<sub>1</sub>2<sub>1</sub>2 | *P*4<sub>1</sub>2<sub>1</sub>2 |
| a,b,c (Å) | 155.67, 155.67, 138.56 | 154.80, 154.80, 135.68 | 156.21, 156.21, 137.82 |
| α, β, γ (°) | 90.00, 90.00, 90.00 | 90.00, 90.00, 90.00 | 90.00, 90.00, 90.00 |
| R<sub>merge</sub><sup>b</sup> | 0.242 (2.587) | 0.430 (7.592) | 0.216 (5.483) |
| R<sub>pim</sub><sup>c</sup> | 0.047 (0.496) | 0.104 (1.797) | 0.042 (1.046) |
| CC<sub>1/2</sub><sup>d</sup> | 0.998 (0.350) | 0.994 (0.309) | 0.999 (0.342) |
| Redundancy | 27.4 (28.0) | 17.5 (17.9) | 27.3 (28.3) |
| Completeness (%) | 100.0 (100.0) | 100.0 (100.0) | 100.0 (100.0) |
| I/σ | 13.0 (1.8) | 6.6 (0.7) | 15.3 (0.9) |
| Refinement |  |  |  |
| Resolution (Å) | 2.10 | 2.66 | 2.56 |
| No. reflections | 98445 (6918) | 47900 (3312) | 55062 (2697) |
| R<sub>work</sub>/R<sub>free</sub><sup>e</sup> | 0.200/0.227<br>(0.306/0.303) | 0.201/0.251<br>(0.356)/(0.415) | 0.228/0.255<br>(0.322/0.335) |
| Number of atoms<sup>f</sup> |  |  |  |
| Protein | 7946 | 7798 | 7812 |
| Ligands | 102 | 88 | 42 |
| Solvent | 256 | 52 | 66 |
| Average B factors (Å<sup>2</sup>) |  |  |  |
| Protein | 40 | 65 | 68 |
| Ligands | 42 | 69 | 73 |
| Solvent | 38 | 58 | 64 |
| RMS deviations |  |  |  |
| Bond lengths (Å) | 0.007 | 0.008 | 0.003 |
| Bond angles (°) | 0.8 | 1.2 | 0.6 |
| Ramachandran plot<sup>g</sup> |  |  |  |
| Favored (%) | 98.10 | 95.98 | 96.86 |
| Allowed (%) | 1.90 | 3.82 | 3.14 |
| Outliers (%) | 0.00 | 0.20 | 0.98 |
| Molprobity score | 1.12 | 1.93 | 1.30 |
| PDB entry | 37SR | 37MV | 38DP |

<sup>a</sup>Values in parentheses refer to the highest-resolution shell of data.
<sup>b</sup>R<sub>merge</sub> = ∑<sub>*h*</sub>∑<sub>*i*</sub>|I<sub>*i,h*</sub> − ⟨I⟩<sub>*h*</sub>|/∑<sub>*h*</sub>∑<sub>*i*</sub>I<sub>*i,h*</sub>, where ⟨I⟩<sub>*h*</sub> is the average intensity calculated for reflection *h* from *i* replicate measurements.
<sup>c</sup>R<sub>p.i.m.</sub> = (∑<sub>*h*</sub>(1/(N-1))<sup>1/2</sup>∑<sub>*i*</sub>|I<sub>*i,h*</sub> −⟨I⟩<sub>*h*</sub>|)/∑<sub>*h*</sub>∑<sub>*i*</sub> I<sub>*i,h*</sub>, where N is the number of reflections and ⟨I⟩<sub>*h*</sub> is the average intensity calculated for reflection *h* from replicate measurements.
<sup>d</sup>Pearson correlation coefficient between random half-datasets.
<sup>e</sup>R<sub>work</sub> = ∑||F<sub>o</sub>| − |F<sub>c</sub>||/∑|F<sub>o</sub>| for reflections contained in the working set. |F<sub>o</sub>| and |F<sub>c</sub>| are the observed and calculated structure factor amplitudes, respectively. R<sub>free</sub> is calculated using the same expression for reflections contained in the test set held aside during refinement.
<sup>f</sup>Per asymmetric unit.
<sup>g</sup>Calculated with MolProbity.

## Results

### Product assays

Incubation of wild-type AsDMS with FPP reveals robust generation of drimenol in overnight (15 h) assays (Figure 2). Incubation of D33A-D323A AsDMS with FPP yields no extractable products, confirming that the double mutant is fully inactivated. Surprisingly, incubation of D33A AsDMS with FPP reveals drimenol generation, but substantially diminished relative to wild-type AsDMS. We attribute this to slow, nonenzymatic Mg^2+^-facilitated solvolysis of DPP in the absence of an active phosphatase domain over the course of 15 h; importantly, no drimenol is detected after 2 h, the timescale of channeling assays (Figure S1). Also notable is the fact that incubation of D323A AsDMS with FPP for 15 h yields small amounts of farnesol, indicating that FPP is a relatively poor substrate for the Mg^2+^-dependent phosphatase domain (Figure 2). Consistent with this result, incubation of D323A AsDMS with FPP in the absence of Mg^2+^ yields no extractable products. Moreover, FPP is stable in Mg^2+^-containing buffer so there is no significant Mg^2+^-dependent solvolysis reaction in the absence of enzyme. Assays run for a shorter period of time (1 h) with D333N AsDMS do not reveal enzyme-catalyzed hydrolysis of FPP,^30^ so the FPP hydrolysis activity observed here with D323A AsDMS is weak at best.

**Figure 2.**
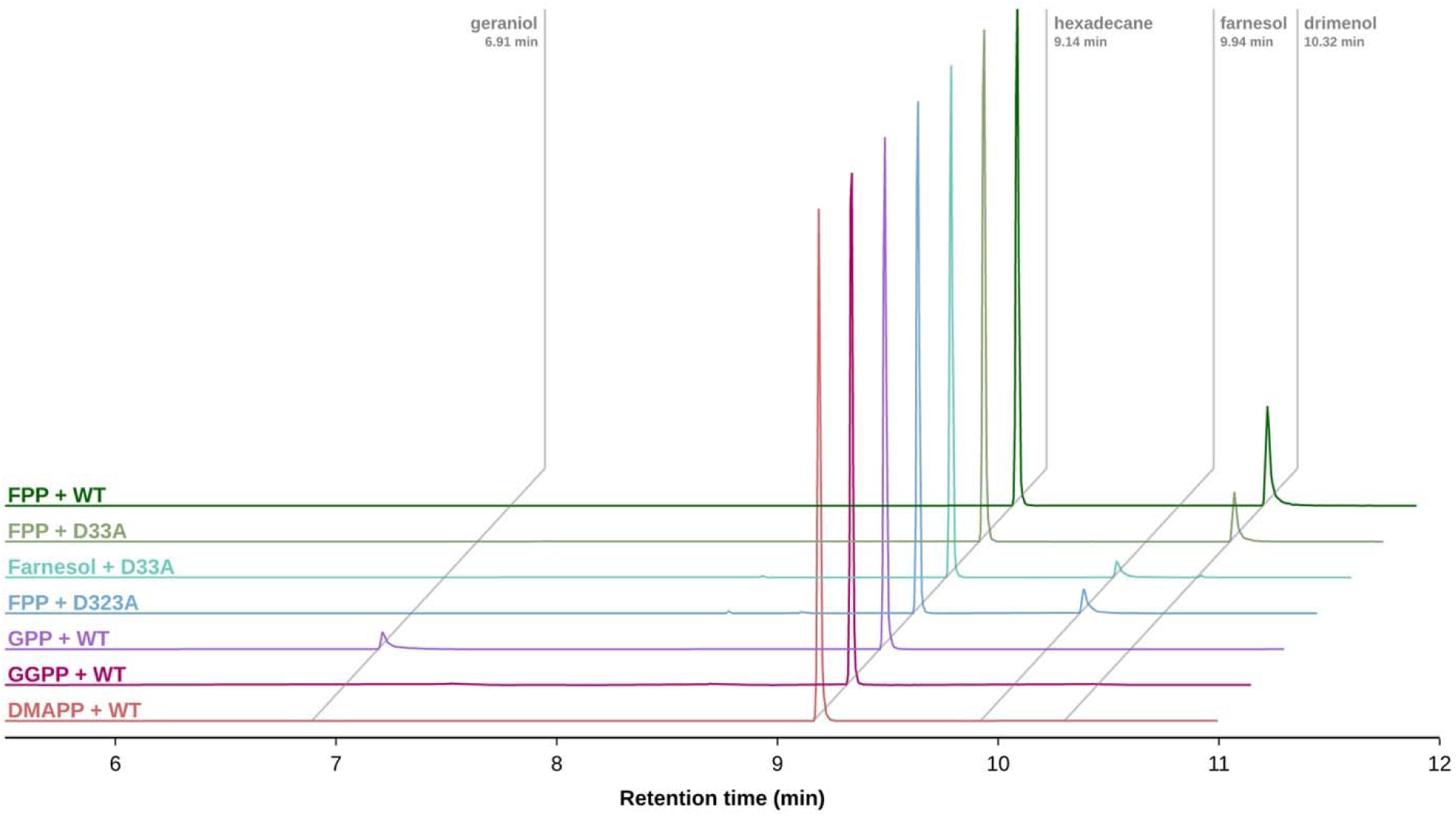
AsDMS product analysis. Gas chromatograms of products formed by wild-type (WT) and mutant AsDMS when incubated overnight (15 h) with various isoprenoids, normalized to a 750 μM hexadecane internal standard in each mixture. Chromatograms for GGPP and DMAPP with an extended retention time axis are found in Figure S1 and similarly indicate no products formed.

Incubation of wild-type AsDMS in overnight assays with GPP reveals generation of geraniol, but incubation with DMAPP or GGPP yields no alcohol product (Figure 2, Figure S1). Thus, weak phosphatase activity appears to be best with medium-sized isoprenoid diphosphates. Interestingly, related HAD-like phosphatases are known to hydrolyze isoprenoid diphosphates such as FPP and GGPP.^39,40^ Finally, incubation of D33A AsDMS with farnesol yields a very tiny but measurable amount of drimenol, suggesting that farnesol is a very poor cyclization substrate relative to FPP.

### Crystal structures

D33A-D323A AsDMS was confirmed to be catalytically inactive and the X-ray crystal structure of its complex with the intact substrate FPP was determined at 2.1 Å resolution. FPP binds in the active site of the cyclase domain with a catalytically productive conformation, i.e., the conformation that would yield the correct regioisomer and stereoisomer of drimenyl diphosphate (Figure 3A). The terminal isoprenoid C=C bond is located approximately 3.9 Å away from A323, poised for the protonation that would initiate catalysis by D323 in the wild-type enzyme. Extended loops containing multiple arginine residues – R368, R370, R415, and R503 – flank the active site, and these residues are partially or fully disordered even upon the binding of non-native ligands.^31^ However, upon the binding of FPP, all arginine residues plus Y416 are fully ordered and donate hydrogen bonds to the substrate diphosphate group (Figure 3A), as previously observed in the D333N AsDMS-FPP complex.^32^

**Figure 3.**
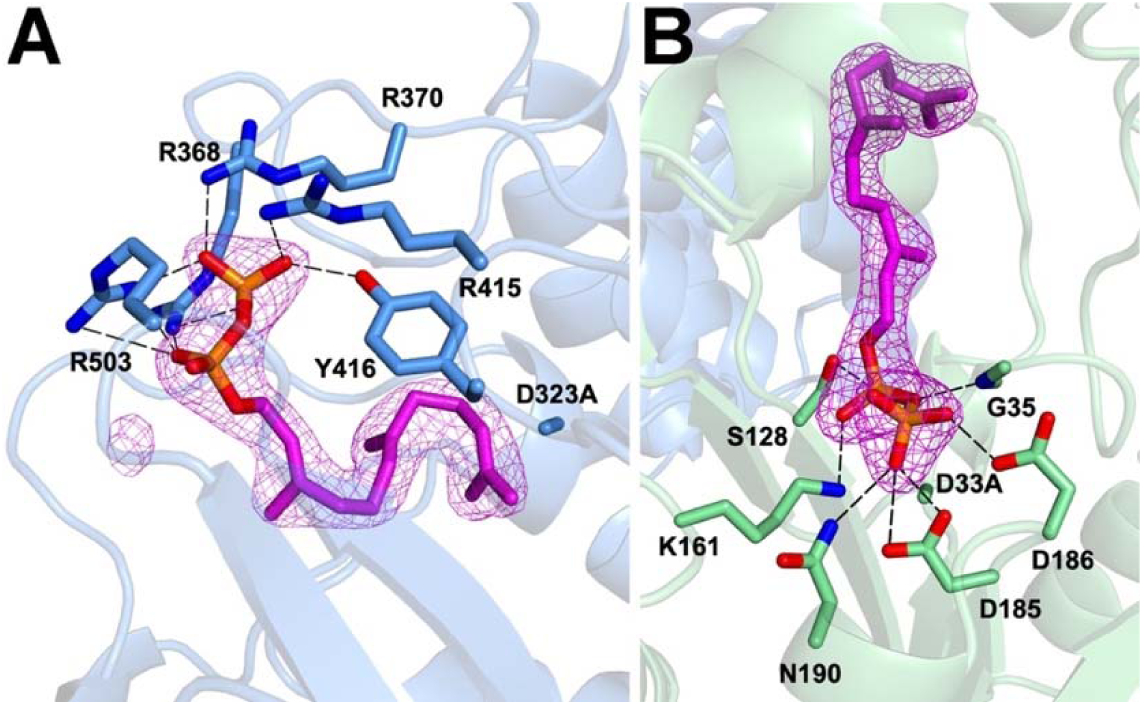
D33A-D323A AsDMS–FPP complex. (A) Polder omit map showing FPP (magenta, contoured at 6.5σ) bound in the active site of the class II cyclase β domain (monomer B). (B) Polder omit map showing FPP (magenta, contoured at 6.5σ) bound in the active site of the HAD-like phosphatase domain (monomer B). Hydrogen bonds are indicated by black dashed lines in each panel and distances are recorded in Table S1.

As also observed in D333N AsDMS,^32^ a second molecule of FPP binds in the phosphatase domain of D33A-D323A AsDMS (Figure 3B). FPP is a poor hydrolysis substrate of the phosphatase domain in wild-type AsDMS (Figure 2), but the D33A mutation abolishes activity by deleting the catalytic nucleophile and metal ligand. The diphosphate group of FPP engages in a network of hydrogen bonds with canonical HAD-like phosphatase motifs,^32,41,42^ including the side chains of S128, K161, D185, D186, N190, and the backbone NH group of G35. The carboxylate side chains of D185 and D186 are presumably protonated to support hydrogen bonding with the diphosphate anion as implied by their close contacts. The isoprenoid moiety of FPP extends into a largely hydrophobic cleft.

Although AsDMS does not catalyze cyclization of the shorter C_10_ isoprenoid GPP, we observed that the phosphatase domain hydrolyzes GPP to yield geraniol (Figure 2). Accordingly, we determined the X-ray crystal structure of the D33A-D323A AsDMS–GPP complex at 2.6 Å resolution to probe structure-activity relationships. The binding conformation of GPP in the cyclase domain resembles that of FPP, and the GPP diphosphate group forms hydrogen bonds with R368, R370, R415, Y416, and R503 (Figure 4A). The isoprenoid tail of GPP curls toward A323 but is 5.8 Å away. With the molecular recognition of GPP dominated by hydrogen bond interactions of the diphosphate group, the terminal isoprenoid C=C bond would be too distant from general acid D323 to enable protonation in the wild-type enzyme. In the phosphatase domain, the diphosphate group of GPP forms hydrogen bonds with the side chains of S128, K161, D185, D186, and N190 (Figure 4B). The side chains of D185 and D186 are likely protonated to accommodate hydrogen bond interactions implied by their close contacts with the diphosphate group. The catalytic Mg^2+^ ion is absent due to the loss of the metal ligand in the D33A mutant.

**Figure 4.**
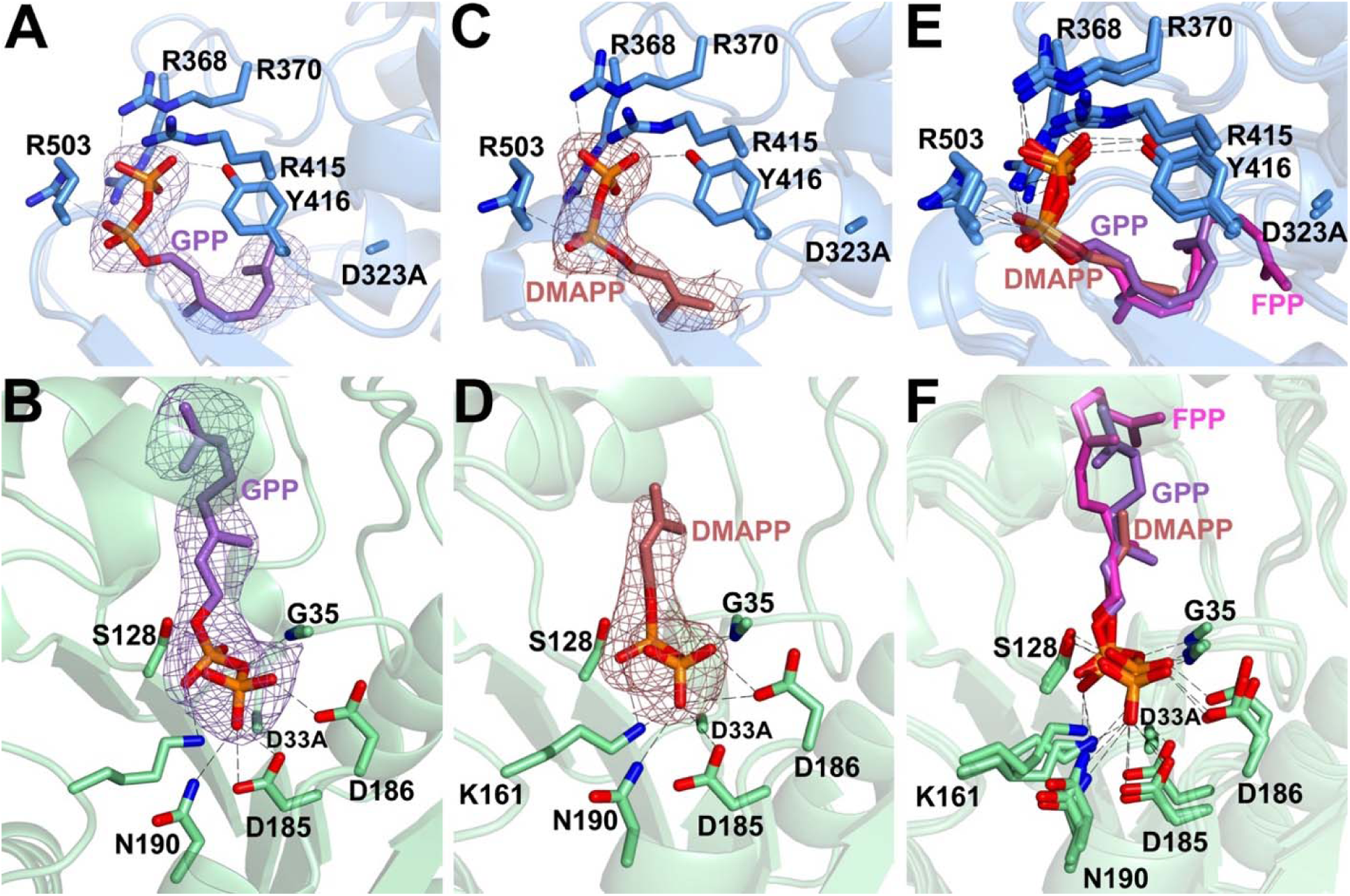
GPP and DMAPP bound to D33A-D323A AsDMS. All maps are contoured at 6.5σ and show monomer B in the asymmetric unit unless otherwise indicated; hydrogen bonds are indicated by dashed lines. (A) Polder omit map showing GPP bound in the active site of the cyclase domain. (B) Polder omit map showing GPP bound in the active site of the phosphatase domain. (C) Polder omit map showing DMAPP bound in the active site of the cyclase domain (monomer A). (D) Polder omit map showing DMAPP bound in the active site of the phosphatase domain. (E) Superposition of the FPP, GPP, and DMAPP complexes with the cyclase domain showing an identical constellation of hydrogen bonds for diphosphate recognition as well as similar isoprenoid conformations. GPP and DMAPP are too short to reach general acid D323 in the wild-type enzyme, so they cannot be substrates. (F) Superposition of the FPP, GPP, and DMAPP complexes with the phosphatase domain showing an identical constellation of hydrogen bonds for diphosphate recognition but divergent isoprenoid conformations. Hydrogen bonds are indicated by black dashed lines in each panel and interatomic distances are recorded in Table S1.

The C_5_ isoprenoid DMAPP is neither a cyclization substrate nor a hydrolysis substrate of AsDMS (Figure 2). Even so, DMAPP binds to both the cyclase and phosphatase domains, as revealed in the 2.56 Å resolution structure of the D33A-D323A AsDMS–DMAPP complex. In the cyclase domain, the DMAPP diphosphate group forms hydrogen bonds with R368, R370, R415, Y416, and R503 with an overall geometry identical to that of the GPP complex; the isoprenoid C=C group is 9.1 Å away from A323 (Figure 4C). In the phosphatase domain, the diphosphate group of DMAPP forms hydrogen bonds with the side chains of S128, K161, D185, D186, and N190 (Figure 4D). Close contacts with D185 and D186 suggest that the side chains of these residues are protonated to accommodate hydrogen bond interactions with the DMAPP diphosphate group. Again, the catalytic Mg^2+^ ion is absent due to the loss of the metal ligand in the D33A mutant.

In both the cyclase domain and the phosphatase domain, the DMAPP diphosphate group engages in a constellation of hydrogen bond interactions identical to that observed in GPP and FPP complexes (Figures 4E,F). With no distinguishing features in the molecular recognition of the diphosphate groups of DMAPP, GPP, and FPP, it is not clear why FPP and GPP are hydrolysis substrates whereas DMAPP is not (Figure 2). Presumably, the structural features responsible for substrate recognition and hydrolysis would be evident upon the binding of the catalytic Mg^2+^ ion to wild-type AsDMS.

### Substrate channeling

One of the key questions remaining with regard to catalysis by AsDMS is whether substrate channeling can occur from the cyclase to the phosphatase. Notably, substrate channeling has been observed in oligomeric bifunctional terpene synthases in which a prenyltransferase is combined with a class I or a class II cyclase.^24,43,44^ In these examples, multiple domains catalyzing sequential reactions are held together in nanoscale oligomeric assemblies thought to enable substrate channeling by proximity. While AsDMS is a monomer in solution at lower concentrations based on initial mass photometry measurements,^31^ mixtures of monomer and dimer are evident in gel filtration chromatography,^32^ and AsDMS crystallizes as a dimer.^31,32^ Here, a more thorough investigation using mass photometry reveals concentration-dependent dimerization (Figure 5, Figure S2).

**Figure 5.**
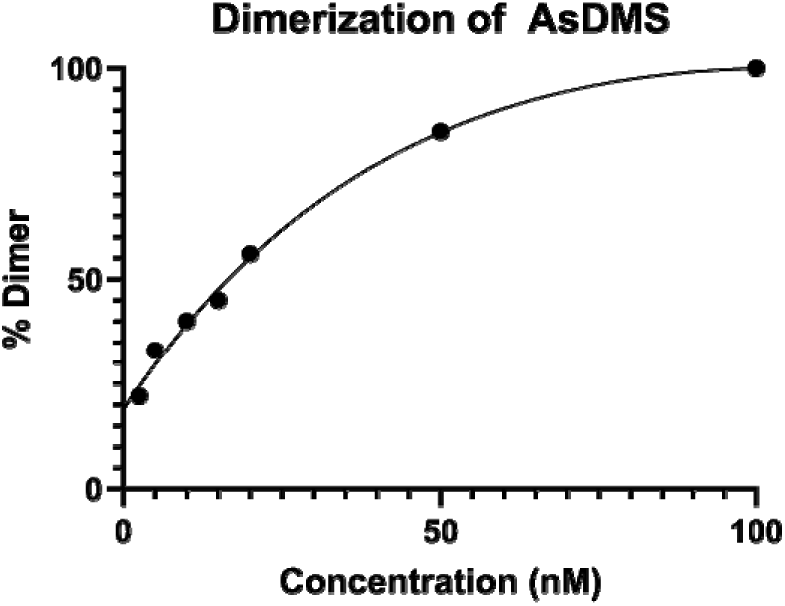
Concentration-dependent dimerization of AsDMS. Individual mass photometry plots are found in Figure S2.

To ascertain the possibility of substrate channeling in AsDMS, we compared the steady-state kinetics of (1) wild-type AsDMS and (2) an equimolar mixture of D33A AsDMS (active cyclase, inactivated phosphatase) and D323A AsDMS (inactivated cyclase, active phosphatase). Since it was not feasible to measure the activities of individual active sites in the wild-type enzyme, we must assume that the cyclase activity of D33A AsDMS and the phosphatase activity of D323A AsDMS are unchanged from activities in the wild-type enzyme. This is a reasonable assumption given that (1) active sites in the monomer are 43 Å apart on opposite sides of the protein (Figure 1), (2) there is no evidence that these active sites communicate with each other, and (3) each mutation is made to a solvent-exposed catalytic residue such that there are no short-range or long-range changes in the surrounding protein structure relative to wild-type AsDMS (Figure S3).

The mixed mutant system was designed to physically uncouple the two catalytic steps, such that the DPP generated by monomeric D33A AsDMS would have to dissociate into bulk solvent and then bind to monomeric D323A AsDMS to generate drimenol. If DPP underwent intramolecular channeling from the cyclase to the phosphatase, such that some or all of this intermediate remained on the enzyme for hydrolysis, wild-type AsDMS should outperform the equimolar mixed mutant system.

However, AsDMS dimerizes at higher concentrations, so the mixed mutant system will form 50% heterodimers and 50% homodimers; heterodimers will contain an active cyclase in one monomer and an active phosphatase in the other. If inter-monomer DPP channeling occurs in wild-type AsDMS, it could still occur in D33A AsDMS-D323A AsDMS heterodimers. Even so, reduced activity would still be observed since heterodimers constitute only 50% of the protein mixture. If intra-monomer DPP channeling occurs in wild-type AsDMS, it could not occur in a homodimer or a heterodimer of the mixed mutant system so reduced activity would be observed. If there is both intramonomer and intermonomer DPP channeling in wild-type AsDMS, reduced activity would still be observed in the mixed mutant system. Thus, reduced activity in the mixed mutant system would provide a qualitative indication of substrate channeling, but not a quantitative measurement of the extent of substrate channeling or whether substrate channeling depends on oligomerization.

Assays were performed at three different AsDMS concentrations: 20 nM (56% dimer), 50 nM (85% dimer), and 100 nM (100% dimer) (Figure 5, Figure S2). Reaction velocities (v_0_) were measured across a range of FPP concentrations for both wild-type AsDMS and the equimolar D33A AsDMS-D323A AsDMS mixture. Across all concentrations of FPP and AsDMS tested, wild-type AsDMS exhibits faster reaction velocities than the equimolar mixed mutant system (Figure 6). Interestingly, the catalytic advantage is greater at lower enzyme concentrations at which the percentage of monomer increases substantially. We conclude that a significant rate enhancement results from covalent linkage of the cyclase and phosphatase domains, consistent with proximity channeling of DPP from the cyclase to the phosphatase.

**Figure 6.**
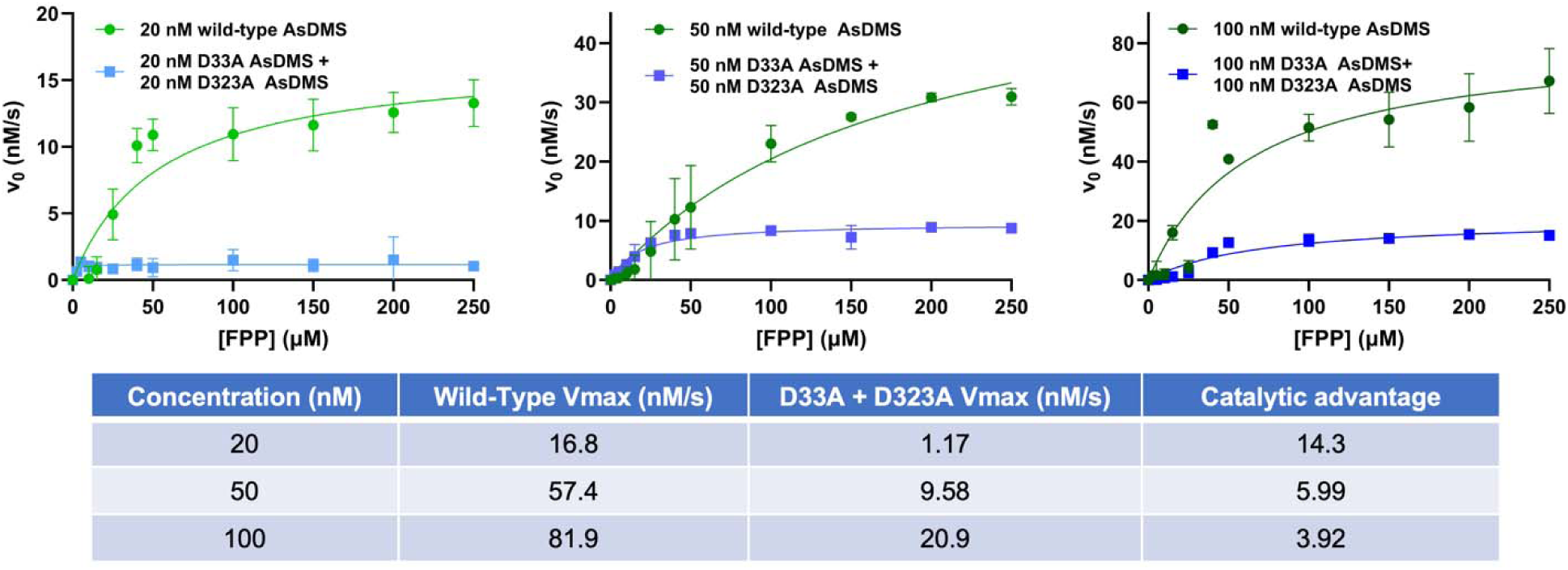
Steady-state kinetics of wild-type AsDMS compared with an equimolar mixture of D33A AsDMS and D323A AsDMS. Initial reaction velocity (v_0_) is plotted as a function of FPP concentration for wild-type AsDMS (green) and an equimolar mixture of cyclase-inactive (D323A) and phosphatase-inactive (D33A) AsDMS variants (blue). Maximum velocity (V_max_) was calculated using a Michalis-Menten regression in GraphPad Prism. Each data point represents the mean of three replicates ± standard deviation.

## Discussion

The active site of a terpene cyclase serves as a template for catalysis, such that the flexible isoprenoid substrate is held in a unique conformation that directs a specific sequence of intramolecular carbon-carbon bond-forming reactions upon initial carbocation formation. The class II cyclase domain of AsDMS clearly serves this function in view of the catalytically productive binding conformations of FPP as first observed in D303A drimenyl diphosphate synthase (SsDMS),^45^ subsequently observed in D333N AsDMS^32^ and D273N AstC,^46^ and here observed in D33A-D323A AsDMS (Figure 7A-C). The crystallographic capture of an intact enzyme-substrate complex in these examples is achieved by substitution of a nonfunctional residue for the aspartic acid general acid that would otherwise initiate the cyclization cascade. Although the conformation of the FPP diphosphate group differs between AsDMS and AstC (Figure 7D), the isoprenoid chain conformations of FPP bound to AsDMS, SsDMS, and AstC are essentially identical and poised to form the correct stereoisomer of drimenyl diphosphate (Figure 7E).

**Figure 7.**
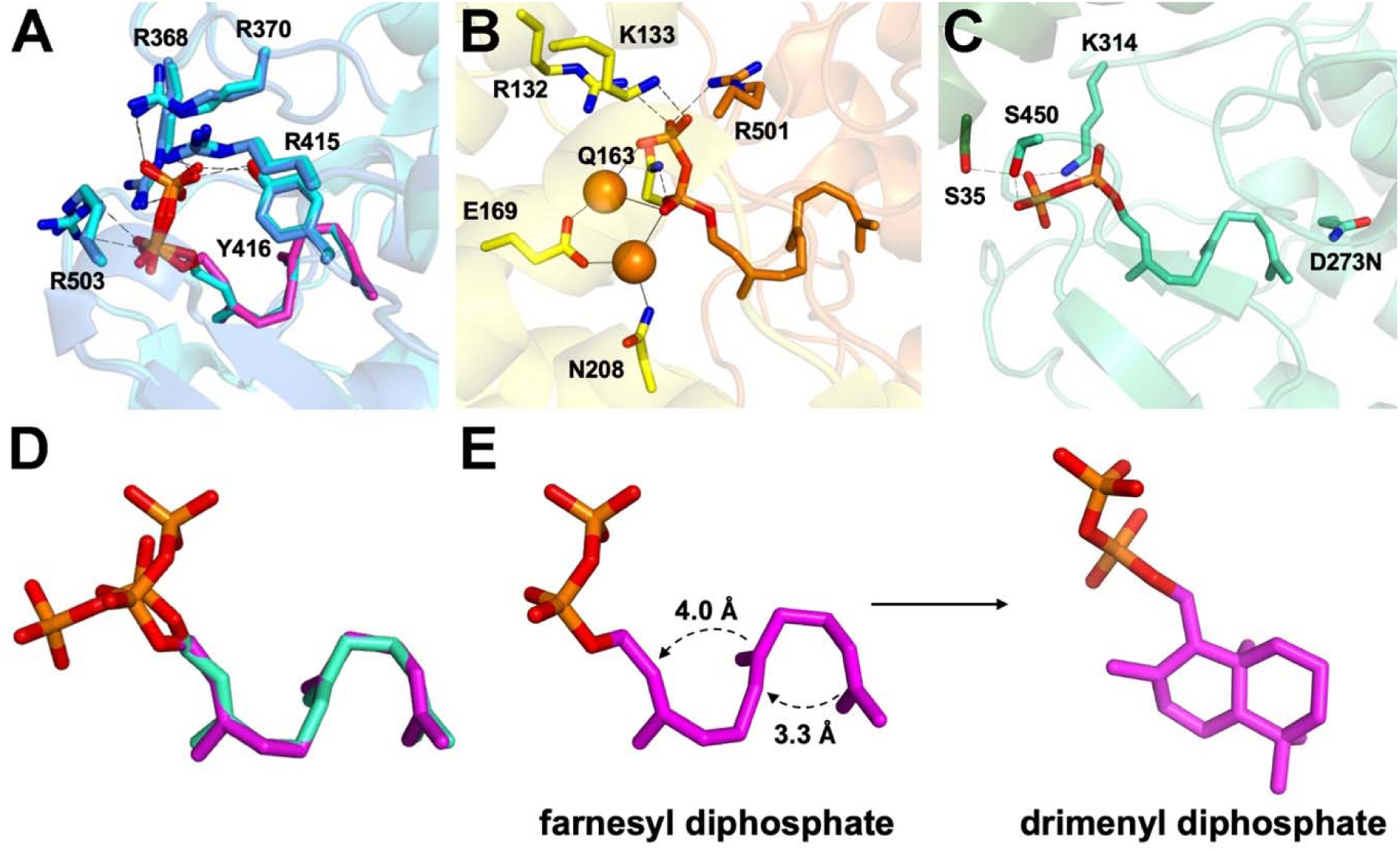
(A) Superposition of FPP complexes with D333N AsDMS (cyan substrate and cyan protein, PDB 9M7F), and D33A-D323A AsDMS (magenta substrate and blue protein, PDB 37SR) reveals essentially identical isoprenoid binding conformations. (B) FPP bound to D303A SsDMS mimics the conformation found in AsDMS, but diphosphate recognition occurs at the interface of the β (orange) and γ (yellow) domains (PDB 7XRA). (C) FPP bound to D273N AstC (PDB 22TZ). (D) Superposition of FPP conformations bound to D33A-D323A AsDMS (magenta) and D273N AstC (green) reveal alternative orientations for the substrate diphosphate group. (E) The common isoprenoid conformation of FPP in (A), (B), and (C) is exactly that required for drimenyl diphosphate generation, as illustrated here for FPP in its complex with D33A-D323A AsDMS. The trajectories of carbon-carbon bond formation are illustrated by dashed curved arrows with carbon-carbon distances indicated.

Most class II cyclases that utilize an isoprenoid diphosphate substrate feature a βγ domain assembly with Mg²⁺-stabilized diphosphate binding, as observed in drimenyl diphosphate synthase (Figure 7B).^45^ Exceptions include class II βγ cyclases that utilize neutral substrates, such as squalene-hopene cyclase, oxidosqualene cyclase, or the meroterpenoid synthase MstB, which do not require Mg^2+^ for substrate binding.^47–49^ AsDMS and AstC are unique among class II cyclases that utilize an isoprenoid diphosphate substrate, not only because these cyclases consist of a single β domain, but also because they do not require Mg^2+^ for substrate binding in the cyclase domain (Figure 7A,C). FPP consistently adopts a conformation poised for drimenyl ring formation independent of its binding to a class II βγ cyclase like SsDMS or a class II β cyclase like AsDMS.

The AsDMS phosphatase complexes with FPP, GPP, and DMAPP show that the HAD-like domain of AsDMS contains conserved S/T and K/D motifs that donate hydrogen bonds to the diphosphate groups of these isoprenoids, yet only FPP and GPP are substrates for hydrolysis. Neither DMAPP nor GGPP are substrates for hydrolysis (Figure S1; the structure of an AsDMS-GGPP complex was not determined). It is likely that binding differences among these isoprenoid diphosphates that govern hydrolysis chemistry in the phosphatase domain would only be discernible in complexes with the catalytically active wild-type enzyme containing an active site Mg^2+^ ion. FPP binding to the phosphatase domain of D333N AsDMS is also observed by Takahashi and colleagues, but Mg^2+^ is not observed to bind even though metal-binding residues are intact in the phosphatase domain (Figure 8).^32^

**Figure 8.**
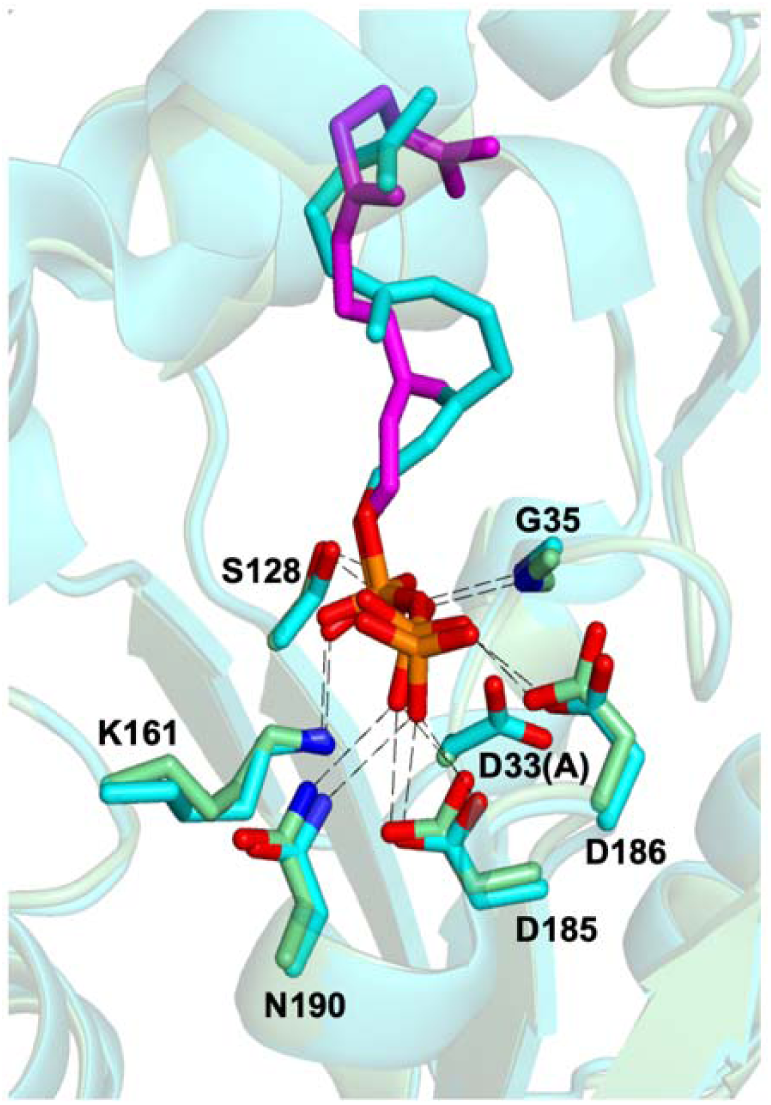
Superposition of the D33A-D323A AsDMS–FPP complex (magenta) and the D333N AsDMS– FPP complex (cyan) showing essentially identical conformations of and hydrogen bond interactions with the diphosphate group of FPP.

Our kinetic evidence for DPP channeling from the cyclase to the phosphatase of AsDMS complements a recent study of a related fungal class II terpene cyclase-phosphatase, albicanoyl monophosphate synthase from *Aspergillus aculeatus* (AacA).^46^ Phosphate-release kinetics measured for dimeric AacA indicate that handoff of the intermediate (albicanoyl diphosphate) from the cyclase to the phosphatase is rate-limiting, rather than intrinsic hydrolysis chemistry. Lin and colleagues suggest that direct transfer of albicanoyl diphosphate from the cyclase of one protomer to the phosphatase of the other is consistent with the observed kinetics.^46^

Takahashi and colleagues suggest that DPP channeling from the cyclase to the phosphatase of AsDMS may be guided by the electrostatic potential of the protein surface, i.e., electrostatic channeling.^32^ Such channeling does not require an enclosed tunnel, but instead requires a route across the protein surface between active sites lined by residues with charges complementary to that of the intermediate in transit. Electrostatic channeling of this nature was first proposed for bifunctional dihydrofolate reductase-thymidylate synthase (DHFR-TS), which contains a cationic surface thought to facilitate transit of the anionic folate intermediate from one active site to the other.^50^ Analysis of the AsDMS structure reveals cationic patches between active sites suggested to serve this function.^32^

It is instructive to compare the functions of AsDMS and AacA, each of which contains class II terpene cyclase and phosphatase domains: AsDMS generates a fully dephosphorylated product, and AacA generates a monophosphate product, albicanoyl monophosphate. That AacA generates a monophosphate product is consistent with our mechanistic proposal for stepwise diphosphate hydrolysis by AsDMS^31^ in which catalytic nucleophile D33 forms two sequential covalent phosphoaspartyl intermediates en route to drimenol. Curiously, the diphosphate hydrolysis reaction catalyzed by AacA stops after one dephosphorylation reaction. Lin and colleagues suggest that the albicanoyl monophosphate product binds too far away from the catalytic aspartate to undergo hydrolysis.^46^

## Concluding remarks

Simultaneous inactivation of both catalytic domains in AsDMS enables crystallographic capture of isoprenoid diphosphate binding in both the class II cyclase domain as well as the phosphatase domain. This approach yields the structure of the precatalytic Michaelis complex in the cyclase domain, with FPP held in the exact conformation required to generate the *trans*-decalin ring system of biosynthetic intermediate DPP. Intriguingly, this approach also yields the structure of FPP bound in the phosphatase domain. Additionally, GPP is a hydrolysis substrate; intact GPP binds in both catalytic domains of D33A-D323A AsDMS. Since the diphosphate groups of FPP, GPP, and DMAPP make essentially identical arrays of hydrogen bond interactions in the phosphatase domain, it is not clear why GPP and FPP are hydrolysis substrates and DMAPP is not. Presumably, substrate binding is significantly influenced by the catalytic Mg^2+^ ion, which is not bound in these complexes due to the loss of metal ligand D33.

Our kinetic evidence indicates that covalent attachment of the class II cyclase domain and the HAD-like phosphatase domain confers a catalytic advantage, such that some of the DPP intermediate remains on the enzyme for transit to the phosphatase domain for hydrolysis. The magnitude of the catalytic advantage is relatively modest and consistent with channeling by proximity.^51^ There is no specific channel or tunnel between the cyclase and phosphatase active sites, but active sites are sufficiently close together such that the DPP intermediate can sometimes dissociate from the cyclase active site and encounter the phosphatase active site before diffusing into bulk solvent. It is not clear to what extent this proximity effect might differ between the AsDMS monomer and dimer, although it is intriguing that the catalytic advantage appears to increase with increasing concentration of the monomer (Figure 6). Future studies of AsDMS as well as other unusual bifunctional chimeras will undoubtedly yield new insight on orthogonal tandem catalysis in terpene biosynthesis.^52^

## ASSOCIATED CONTENT

### Supporting Information

Interatomic distances in AsDMS complexes (Table S1), gas chromatograms (Figure S1), mass photometry results (Figure S2), and wild-type and double mutant structural alignment (Figure S3).

### Accession Codes

The atomic coordinates and crystallographic structure factor amplitudes of the D33A-D323A AsDMS complexes with FPP, GPP, and DMAPP have been deposited in the Protein Data Bank (www.rcsb.org) with accession codes 37SR, 37MV, and 38DP, respectively.

## AUTHOR INFORMATION

### Corresponding Author

**David W. Christianson –** Roy and Diana Vagelos Laboratories, Department of Chemistry, University of Pennsylvania, Philadelphia, Pennsylvania 19104-6323, United States;

### Authors

**Kristin R. Osika –** Roy and Diana Vagelos Laboratories, Department of Chemistry, University of Pennsylvania, Philadelphia, Pennsylvania 19104-6323, United States.

**Mason E. Leffler –** Roy and Diana Vagelos Laboratories, Department of Chemistry, University of Pennsylvania, Philadelphia, Pennsylvania 19104-6323, United States.

**Briana Abigail R. Czarnecki** – Roy and Diana Vagelos Laboratories, Department of Chemistry, University of Pennsylvania, Philadelphia, Pennsylvania, 19104-6323, United States.

## ACKNOWLEDGMENTS

We thank Drs. Matthew Gaynes and Kushol Gupta for many helpful scientific discussions. This work is based on research conducted at beamline 17-ID-2 (FMX) of the National Synchrotron Light Source II, a DOE Office of Science User Facility operated for the DOE Office of Science by Brookhaven National Laboratory under Contract DE-SC0012704. The Center for BioMolecular Structure (CBMS) is primarily supported by the National Institutes of Health, NIGMS, through a Center Core P30 Grant (P30GM133893) and by the DOE Office of Biological and Environmental Research (KP1605010).

## For Table of Contents Use Only

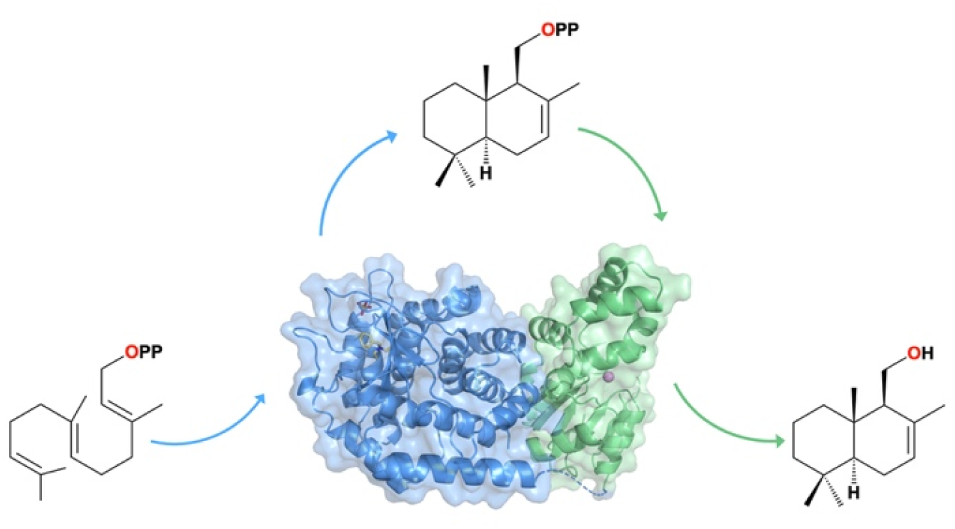

