## Supporting Information for "Isoprenoid Binding and Substrate Channeling in Drimenol Synthase, a Bifunctional Class II Terpene Cyclase-Phosphatase"

### Table S1

| | | Interatomic Distances in D33A-D323A AsDMS-Isoprenoid Complexes Indicated in Figures 3 and 4 (Å) | | | | | | | | | | |
|---|---|---|---|---|---|---|---|---|---|---|---|---|
| | | Cyclase | | | | | Phosphatase | | | | | |
| | | R368 | R370 | R415 | Y416 | R503 | G35 (NH) | S128 | K161 | D185 | D186 | N190 |
| FPP | O15 | | | | | | | 2.8 | | | | |
| | O16 | 3.0 | | | | 2.5, 3.5 | | | 2.6 | | | |
| | O17 | | | | | | 3.1 | | | | | |
| | O19 | 2.8 | | | | | | | | | | |
| | O20 | | | 3.0 | 2.7 | | | | | | 2.4 | |
| | O21 | | 2.8 | | | 2.5 | | | | 2.5, 3.1 | | 3.2 |
| GPP | O13 | | | | | | | 2.9 | | | | |
| | O14 | 3.0 | | | | 2.8 | | | 2.7 | | | |
| | O15 | | | | | | 3.0 | | | | | |
| | O17 | 3.0 | | | | | | | | | | |
| | O18 | | | 3.1 | 2.7 | | | | | | 2.6 | |
| | O19 | | 2.8 | | | 3.5 | | | | 2.7, 3.1 | | 3.1 |
| DMAPP | O7 | 3.1 | | | | 2.9 | | | 3.0 | | | |
| | O8 | | | | | | 3.3 | 2.9 | | | | |
| | O9 | | | | | | 3.0 | | | | | |
| | O11 | 2.9 | | | | | | | | | 2.7 | |
| | O12 | | | 2.7 | 2.6 | | | | | 2.7 | 3.2 | |
| | O13 | | 2.8 | | | | | | 3.2 | | | 3.3 |

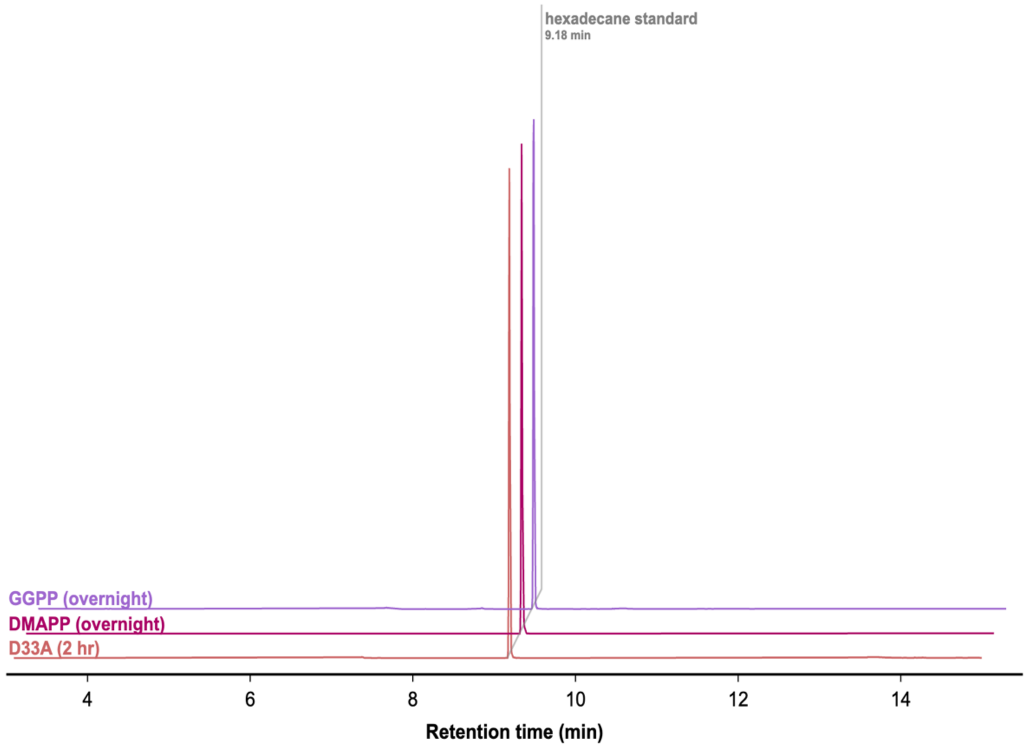


**Figure S1.** Gas chromatograms showing that no extractable products are generated after overnight incubation (15 h) of wild-type AsDMS with GGPP or DMAPP. These chromatograms are identical to those shown in Figure 2 except that the retention time axis extends from 3–15 min. Additionally, no extractable products are generated after incubation of D33A AsDMS with FPP for 2 h; only a 15-h incubation incubation yields a measurable amount of drimenol, as shown in Figure 2.

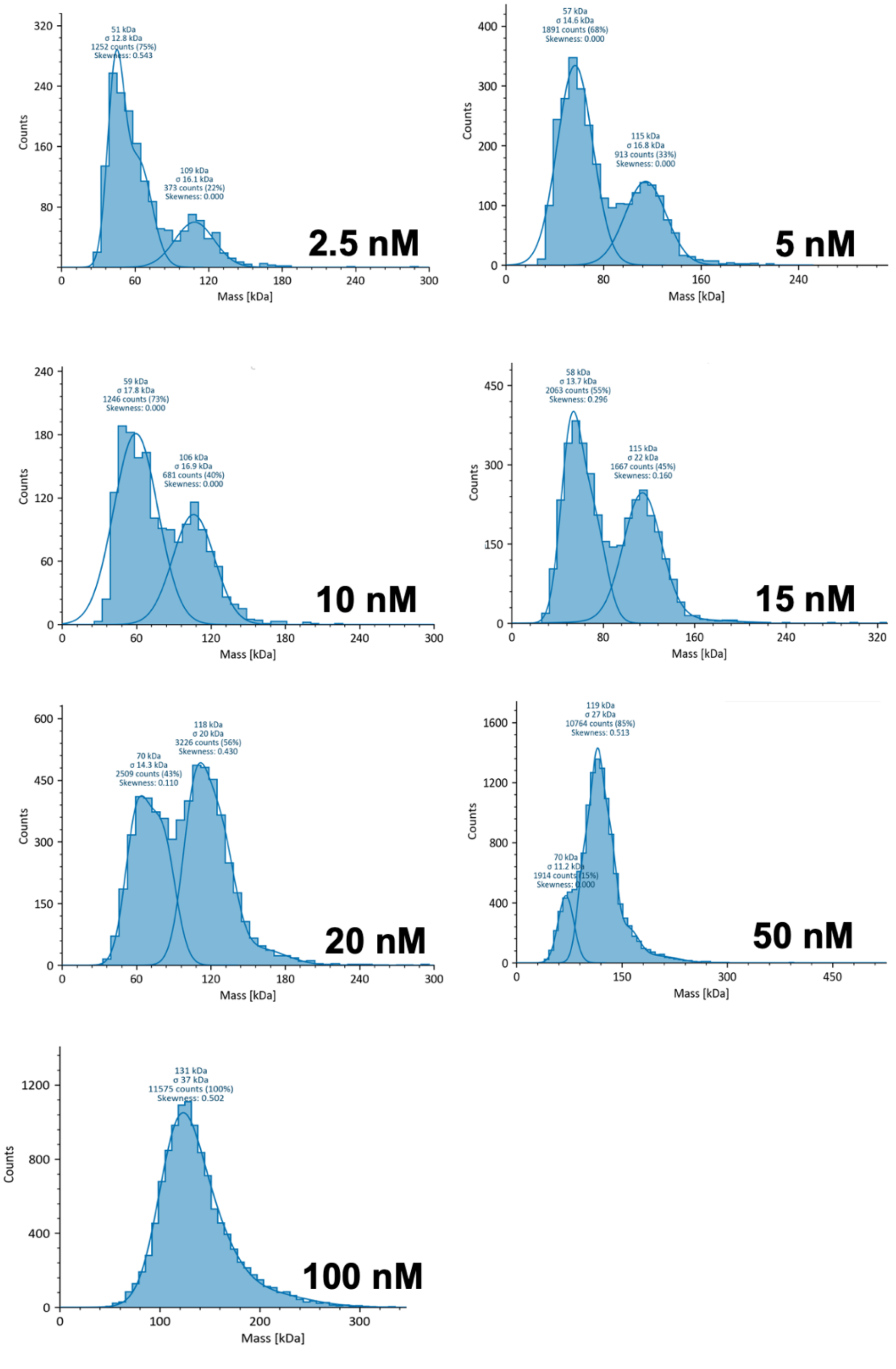


**Figure S2.** Mass photometry measurements indicate that dimerization of AsDMS is concentration-dependent. The molecular weight of the N-terminal deletion variant prepared for this study (residues 1–526) is 61 kD (dimer 122 kD); species observed range from 51-70 kD (monomer) and 106-131 kD (dimer).

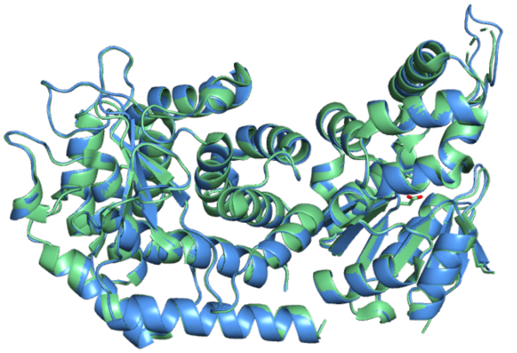

**Figure S3.** Alignment of wild-type AsDMS (green, PDB 9MHS) and double mutant D33A-D323A AsDMS (blue, PDB 37SR). No structural changes result from mutation of catalytic aspartate residues, either within or across their respective catalytic domains.
